# Polo-like kinase 2 inhibition reduces serine-129 phosphorylation of physiological nuclear alpha-synuclein but not of the aggregated alpha-synuclein

**DOI:** 10.1101/2021.05.21.445104

**Authors:** Sara Elfarrash, Nanna Møller Jensen, Nelson Ferreira, Sissel Ida Schmidt, Emil Gregersen, Marie Vibeke Vestergaard, Sadegh Nabavi, Morten Meyer, Poul Henning Jensen

## Abstract

Accumulation of aggregated alpha-synuclein (α-syn) is believed to play a pivotal role in the pathophysiology of Parkinson’s disease (PD) and other synucleinopathies. α-Syn is a key constituent protein of Lewy pathology, and α-syn phosphorylated at serine-129 (pS129) constitutes more than 90% of α-syn in Lewy bodies and hence, it is used extensively as a pathological marker for the aggregated form of α-syn. However, the exact role of pS129 remains controversial as well as the kinase(s) responsible for the phosphorylation.

In this study, we investigated the effect of Polo-like kinase 2 (PLK2) inhibition on formation of pS129 using ex-vivo organotypic brain slice model of synucleinopathy. Our data demonstrated that PLK2 inhibition has no effect on α-syn aggregation, pS129 or inter-neuronal spreading of the aggregated α-syn seen in the organotypic slices. Instead, PLK2 inhibition reduced the soluble nuclear pS129 level confined in the nuclei. The same finding was replicated in an in-vivo mouse models of templated α-syn aggregation and human dopaminergic neurons, suggesting that PLK2 is more likely to be involved in S129 phosphorylation of soluble non-pathology related fraction of α-syn. We also demonstrated that reduction of nuclear pS129 but not the aggregates specific pS129 following PLK2 inhibition for a short time before sample collection improves the signal to noise ratio when quantifying pS129 aggregate pathology.

## Introduction

Mounting evidence from biochemical, pathological and genetic studies strongly suggest a role of alpha-synuclein (α-syn) in the pathogenesis of a group of neurodegenerative diseases, collectively called synucleinopathies [1]. These diseases, which include Parkinson’s disease (PD), dementia with Lewy bodies (DLB) and multiple system atrophy (MSA), share a common pathological hallmark, namely the development of inclusions containing α-syn aggregates in affected brain cells. The aggregated α-syn is a main constituent of Lewy pathology detected in neuronal axons and soma in PD and DLB, and in oligodendrocytes as glial cytoplasmic inclusions in MSA [1]. α-Syn is subject to multiple types of posttranslational modifications [2], and phosphorylation at serine-129 (pS129) has attracted special attention as approximately 90% of aggregated α-syn in the brains of PD patients is stably phosphorylated at this site [3], [4], [5]. In contrast, physiological α-syn is transiently phosphorylated on S129 with less than 4% phosphorylated [3], [4].

Despite the advantages of pS129 as a biomarker for pathological α-syn aggregates, its pathophysiological role has been contested [6], [7], [8], [9], [10]. Some studies demonstrate an increased toxicity associated to S129-phosphorylation [8], [9], [11], whereas others suggests it has no role in α-syn aggregation or toxicity [12], [13] or even being a protective modification [6], [7].

To date, in vitro and in vivo studies have determined a number of kinases able to phosphorylate α-syn at S129, including the Polo-like kinases (PLK) 2 and 3, casein kinases 1 and 2, Leucine-rich repeat kinase 2 (LRRK2) and various G-protein coupled receptor kinases (GRKs) [14], [11], [15], [16], [17], [18]. Of these, PLK2 has been extensively studied [19], [20], [21], [10], [22] but mostly in relation to phosphorylation of physiological, non-aggregated α-syn or the regulation of α-syn expression [19], [21], [22], [23].

In this study, we used a validated PLK2-inhibitor, compound 37 [24], [21], [23], and the PLK1-3 inhibitor BI2536 [25], to investigate the role of PLK2 in the S129-phosphorylation of α-syn aggregates in PD models induced by pre-formed α-syn fibrils (PFFs).

We employed multiple model systems of α-syn aggregation, including mouse organotypic brain slices, α-syn-transgenic mice, and human dopaminergic cell cultures to assess the potential of inhibiting PLK2 on development of pS129-positive aggregates or dephosphorylation of such already formed. We demonstrate that inhibition of PLK2 could neither revert nor prevent S129-phosphorylation of α-syn aggregate inclusions or prevent the spreading of α-syn aggregate pathology between neurons. In contrast, PLK2 was responsible for a significant part of the basal α-syn S129-phosphorylation predominantly located in neuronal nuclei.

## Materials and Methods

### 1. Preparation of organotypic hippocampal slice cultures (OHSCs)

Organotypic hippocampal slices were prepared from 5-7-day-old C57Bl6 pups according to Stoppini et al. 1991 [26] with slight modifications as described in [13]. Briefly, the hippocampi were extracted in carbonated low Na cerebrospinal fluid (CSF) (1 mM CaCl_2_, 10 mM D-glucose, 4 mM KCl, 5 mM MgCl_2_, 26 mM NaHCO_3_, 234 mM sucrose and 0.1% phenol red solution) and coronal slices of 400 μm were made using a tissue chopper (Stoelting, #51425). Hippocampal slices with intact dentate gyrus (DG) and cornu ammonis (CA) regions were selected and maintained on air-fluid interface-style Millicell culture inserts, 30 mm diameter, 0.4 μm (Millipore) in 6-well culture plates (ThermoFisher Scientific) with 800 μL of 37°C pre-heated sterile medium (MEM Eagle medium 78.8% (Gibco #11095), 20% heat-inactivated horse serum (Gibco, #16050-122), 1 mM L-glutamine, 1 mM CaCl_2_, 2 mM MgSO_4_, 170 nM insulin, 0.0012% ascorbic acid, 12.9 mM D-glucose, 5.2 mM NaHCO_3_, 300 mM Hepes (Sigma #H3375), pH=7.28, osmolality adjusted to 317-322). The medium was replaced completely three times per week.

### 2. Microinjection of OHSCs with S129A PFFs

PFFs were produced from monomeric human recombinant α-syn with residue serine-129 shifted to alanine (S129A). S129A PFF production, characterization and validation of efficient aggregation was done as described in [13]. The usage of S129A PFFs that are non-phosphorylatable at S129 ensures that only the endogenously expressed α-syn is detected when antibodies against pS129 are used during analysis.

Immediately before injection, an aliquot of S129A PFFs was thawed at room temperature (RT) and sonicated for 30 seconds using Branson Sonifier 250 with settings adjusted to 30% duty cycle, output control 3. The sonicator is customized and equipped with a water jacket cooling system to avoid sample heating during sonication. OHSCs were microinjected with S129A PFFs or sterile phosphate buffered saline (PBS) at DG after 7 days in culture. Microinjection pipettes (item #1B200F-4 (with Filament), WPI) were pulled using a micropipette puller (P-1000, Sutter Instrument). For injection, a Pulse Pal v2 (#1102) was set to phase 1 voltage 5V, phase 1 duration 0.01 seconds, pulse interval 0.5 seconds. The pipette was loaded using Eppendorf microloader pipette tips (ThermoFisher). A final volume of 0.1 μL of either S129A PFFs (1 mg/mL) or PBS was injected at DG under microscopic guidance as described in [13].

Injections were performed under strict aseptic condition in a laminar flow hood equipped with a microscope. After injecting all slices on a culture insert, the medium was replaced with fresh pre-heated medium.

### 3. Polo-like kinase 2 (PLK2) inhibition in OHSCs

After 6 days in culture, 24 hours before S129A PFF microinjection, organotypic hippocampal slices were treated with either 20 μM PLK2i (compound 37 custom synthesized by Wuxi AppTec, Shanghai, China) [21], [23], or 0.2% DMSO (Sigma, #RNBF6889) as a solvent control. The drug activity and specificity of PLK2i were validated earlier in our lab as described earlier in [23]. Medium containing the drug/vehicle was added below the membrane insert along with a drop of 1 μL on the slice surface to facilitate an equal distribution of the drug throughout the organotypic brain slice thickness [27]. The drug/vehicle mixture was added throughout the experiment with each change of the medium, three times per week.

To evaluate the effect of short-term PLK2 inhibition on PFF-seeded OHSCs, organotypic slices were treated as described above with either 20 μM PLK2i (compound 37) or 1 μM of the PLK1-3 inhibitor BI2536 (Selleck, #S1109) [17] for 24 hours before fixation.

### 4. Immunoblotting and sequential biochemical fractionation of OHSCs

7 days following PFF injection, slices were cut from the culture membrane, maintaining a small border of membrane around the slice to ensure that all the tissue was collected. For each group, ten slices were analyzed. Protein quantification, sequential fractionation into RIPA-soluble and - insoluble fractions and subsequent immunoblotting was carried out as described in [13]. Antibodies used were: rabbit polyclonal anti-α-syn (ASY-1, 1:1000) [28], mouse monoclonal pS129-α-syn (11A5, kindly provided by Imago Pharmaceuticals, 1:2000) [4], rabbit monoclonal rodent-specific α-syn (D37A6, #4179, Cell Signaling, 1:1000) and mouse monoclonal anti-β-tubulin III (TUJ1, #T8578, Sigma, 1:5000). PageRuler pre-stained protein ladder 10-180 kDa (ThermoFisher, #26616) was used as the molecular size marker.

### 5. Immunofluorescence staining of OHSCs

7 days following S129A PFF-injection, OHSCs were fixed using 4% PFA in PBS (2.8 mM NaH_2_PO_4_·H_2_O, 7.2 mM Na_2_HPO_4_·2H_2_O, 123 mM NaCl, pH adjusted to 7.2) and processed for immunohistochemistry as described in [13]. Antibodies used were α-syn aggregate-specific antibody MJF-14 (rabbit monoclonal MJFR-14-6-4-2, #ab209538, Abcam, 1:25,000) and mouse monoclonal pS129-α-syn (11A5, kindly provided by Imago Pharmaceuticals, 1:25,000) [4]. After washing off unbound primary antibody, slices were incubated with the appropriate Alexa Fluor-labelled (488 and 568) secondary antibodies (Invitrogen, 1:2000) and 4’,6-diamidino-2-phenylindole (DAPI) (TH.GEYER, 5 μg/mL) in 5% BSA/PBS.

### 6. M83 mouse treatment with PLK2i

Animals were housed in a temperature-controlled room, under a 12 h light/dark period, with water and food ad libitum. Twelve-month-old M83^+/-^ α-syn transgenic mice were anaesthetized by 3.5% isoflurane inhalation and bilaterally injected with recombinant mouse α-syn PFFs by inserting the needle ~1 mm deep into the biceps femoris as described by [29]. Recombinant mouse α-syn PFFs were prepared and validated for aggregation properties as described previously for human α-syn PFFs in [13] (Supplementary fig. 1). Injections were made using a 10-μL Hamilton syringe with a 25-gauge needle.

Once mice displayed hind limbs paralysis (typically 10-12 weeks post-injection) mice were treated for 2 days with PLK2i (Lundbeck, compound 37) by oral gavage (n=5, 2×100 mg/kg/day) or vehicle control (5% DMSO, 1% methylcellulose, n=5). Then, mice were anaesthetized and perfused with PBS with 1× complete protease inhibitor cocktail (cOmplete, Roche) and phosphatase inhibitors (25 mM β-glycerolphosphate, 5 mM NaF, 1 mM Na_3_VO_4_, 10 mM Na-pyrophosphate), followed by 4% PFA/PBS. Brains were removed and kept in 4% PFA/PBS for 48 hours at 4°C, and then stored in 30% sucrose in PBS with 0.05% NaN_3_ until cryosectioning.

### 7. Cryosectioning and immunofluorescence staining of M83 mouse brain

For cryosectioning, brains were mounted on the cryostate stage (Leica) using Tissue-Tek® O.C.T. Compound (Sakura). After the OCT solidified, the brain was sliced at a thickness of 10-12 μm at −20°C. Sections were collected on Superfrost Plus Adhesion Microscope Slides (ThermoFisher) and subsequently processed for immunostaining. Tissue was permeabilized in 0.5% Triton X-100 followed by blocking with 10% BSA, for 45 minutes each at room temperature (RT) with gentle shaking. For primary antibody incubation, rabbit monoclonal pS129 antibody (D1R1R, #23706, Cell Signaling Technology, 1:1000) was prepared in 5% BSA, and incubated overnight at 4°C. Slides were washed 3x 20 minutes in TBS + 0.03%Triton X-100, and then incubated with anti-rabbit Alexa Fluor 488 (Invitrogen, 1:2000) and DAPI (TH.GEYER, 5 μg/mL) for nuclear staining in 5% BSA for 2 hours at RT, protected from light. During antibody incubation, slides were kept in a humidity chamber with a hydrophobic barrier around the tissue to prevent them from the drying out. After the final washing step, the slides were mounted using DAKO fluorescent mounting medium (DAKO, S3023). The edges of the coverslip were sealed using nail polish.

### 8. Neural stem cell differentiation, treatment and immunostaining

Healthy, human induced pluripotent stem cell-derived neural stem cells were propagated and differentiated using Induction/DOPA Differentiation kit (XCell Science), as previously described [30]. At day 38 of differentiation, neurons were exposed to human recombinant α-syn S129A PFFs (14 μg/mL) added to the cell culture medium. After 24 hours, cells were carefully washed in PBS, and allowed to grow for an additional 6 days in fresh medium before fixation in 4% PFA at day 45. 4 hours prior to fixation, cells were treated with 1 μM BI2536 for inhibition of PLK1-3 [17] or DMSO as a vehicle control. Immunostaining was carried out as described in [23], using the following primary antibodies: chicken polyclonal MAP2 (#ab92434, Abcam, 1:2000), mouse monoclonal pS129 (11A5, 1:10,000) and tyrosine hydroxylase (TH, #AB152, Merck Millipore, 1:1000). Appropriate secondary Alexa Flour antibodies (Life Technologies and Abcam) were diluted 1:1000 and DAPI (TH.GEYER, 5 μg/mL) was used for nuclear staining. Coverslips were mounted with DAKO fluorescent mounting medium (DAKO, S3023) and edges sealed with nail polish.

### 9. Primary neuronal cultures, treatment and immunostaining

Primary hippocampal neurons were prepared from wild type P0 C57Bl6 pups, as previously described [31]. Briefly, hippocampi were dissected in ice-cold PBS and dissociated using papain for 20 minutes at 37°C. Hippocampi were then triturated in plating medium (MEM, Gibco, #51200-020) supplemented with 10% heat-inactivated fetal bovine serum (FBS), 0.5% w/v glucose, 2.38 mM NaHCO_3_, 1.3 μM transferrin (Calbiochem, #616420), 20 mM Glutamax, 86.2 μM insulin (Sigma, #I6634)) and seeded on Matrigel matrix-coated coverslips (Corning, #354234). After 24 hours, medium was changed to growth medium (MEM, Gibco, #51200-020) supplemented with 5% FBS, 0.5% w/v glucose, 2.38 mM NaHCO_3_, 1.3 μM transferrin (Calbiochem, #616420), 5 mM Glutamax, 1x B-27 supplement), and at 3 days in-vitro (DIV) glial proliferation was inhibited with 2 μM cytosine arabinoside (Sigma, C6645).

At 14 DIV, neurons were treated with the PLK1-3 inhibitor BI2536 (10 nM) for 0-4 hours before fixation in 4% PFA and processing for immunocytochemistry as previously described [23]. Primary antibodies used were pS129 (D1R1R, #23706, Cell Signaling Technology, 1:1000) and chicken polyclonal MAP2 (#ab92434, Abcam, 1:1000). Appropriate secondary antibodies (Alexa Fluor, Life Technologies and Abcam) were diluted 1:1000, and DAPI (TH.GEYER, 5 μg/mL) was added for nuclear staining. Coverslips were mounted with DAKO fluorescent mounting medium (DAKO, S3023) and edges sealed with nail polish.

### 10. Quantification

Western blot band density was calculated using ImageJ (National Institutes of Health) after first assuring that the bands were not saturated. Background was subtracted and the density of each band normalized to the density of the loading control (β*III*-tubulin). Quantifications display the mean of three independent experiments. For immunostainings of organotypic slices (Fig. 2–4), four images covering the whole slice were taken using the X10 objective on a Zeiss AxioObserver 7 inverted fluorescence microscope fitted with an ApoTome to increase z plane resolution and analyzed in ImageJ. The focus of each image was adjusted to maximize the amount of visible aggregate pathology. For each image, tissue area was approximated by selection of DAPI-staining of the nuclei and expanded 25 pixel units. Aggregates were defined by the MJFR-14-6-4-2 staining; for each image, uneven background was subtracted by the rolling ball algorithm (size = 25 pixels), the image was thresholded using the Auto Threshold plugin (method = RenyiEntropy), and particles with a minimum size of 6 pixels^2^ were counted as aggregates. For comparisons of aggregate areas, the total aggregate area for each organotypic slice was normalized to its tissue area. For analysis of fluorescent intensities, the mean fluorescent intensity (MFI) of either the MJF-14-staining or the pS129-staining was measured inside the MJFR-14-6-4-2-based definition of aggregates (defined as above) or outside the aggregate definition but inside tissue area. The pS129 fraction defined as “nuclear pS129” when it colocalizes with DAPI staining. The MFI ratio, defined as pS129 MFI inside aggregates divided by pS129 MFI outside aggregates, was used to compare the “signal-to-noise” ratio of the pS129-staining before and after PLK2 inhibition.

For evaluation of a possible effect of the PLK2i-treatment on spreading of aggregation throughout the hippocampal slices (Fig. 3), the previous images were stitched together using the Stitching plugin in Fiji (Fiji Is Just ImageJ, NIH) [32], and the DG and CA1 regions were defined manually from the DAPI-staining (Fig. 3a). First, the DG was bound by a rectangle, and second, projections along one side of the rectangle and through one diagonal were used to define the limits of the CA1 region. The subsequent definition of aggregates and analysis of aggregate areas was performed as described above, based on the MJFR-14-6-4-2 staining.

For M83 mouse brain, 2-5 images per brain region were taken randomly with an X10 (2 images, region at frontal cortex), an X40 oil (3 images, region at DG and CA1 of the hippocampus) or an X63 oil objective (5 images, region at hind brain). For the regions hippocampus (both DG and CA1) and frontal cortex (Fig. 5), only baseline nuclear pS129 was assessed, for hindbrain region, both the nuclear pS129 and the aggregates were both quantified. For analysis, DAPI-staining was used to define nuclei, and the MFI of nuclear pS129 was computed inside this DAPI-selection. For quantification of images from the hind brain (where aggregate pathology was present, Supplementary fig. 3), pS129-positive aggregates were defined by the Auto Threshold plugin (method = MaxEntropy), with a minimum particle size of 200 pixels^2^. pS129 aggregate MFI was then measured inside this selection, and pS129 non-aggregate MFI outside this aggregate selection. Again, the MFI ratio (aggregate-related pS129 divided by non-aggregate pS129) was computed. For human dopaminergic neurons (Fig. 6), 5 X20 images were taken randomly per coverslip. For each image, the number of nuclei was counted in Fiji using the built-in watershed algorithm, and the MAP2+ cell area was defined using the Auto Threshold plugin (method = default). pS129-positive aggregates (inside the MAP2+ cell area) were defined as following: uneven background was subtracted using the rolling ball algorithm (size = 50 pixels), and aggregates were defined as signals with a fluorescent intensity minimum 14x above median intensity (of the entire image) and with a minimum size of 10 pixels^2^. MFI of non-aggregate pS129 was computed as pS129 intensity outside of aggregates (defined as above), but inside the neuronal cell area. Total aggregate area per image was normalized to the number of nuclei. A minimum of 2000 cells was analyzed per condition for each replicate, and two replicates formed the basis for the quantifications.

For primary hippocampal neurons (Supplementary fig. 4), 10-15 X20 images were taken per coverslip, selected based on MAP2-staining to ensure the presence of neurons. Neuronal nuclei were defined by the DAPI-staining co-localized with MAP2-staining, and pS129 MFI inside the neuronal nuclei was computed in ImageJ.

### 11. Statistical analysis

Data were tested for normality using the Shapiro-Wilks test, and normally distributed data were compared using a two-tailed Student’s T-test for comparison of two groups or Welch’s T-test when comparing groups with unequal variances. Non-normally distributed data were compared using the non-parametric two-tailed Mann-Whitney U test. For comparison of multiple groups, one-way ANOVA was conducted, followed by the Holm-Šidák post hoc test or Fisher’s LSD (least significant difference). Data are presented as means ± standard deviation (SD) except where otherwise mentioned. A p-value below 0.05 was considered significant. *p<0.05, **p<0.01, ***p<0.001.

## Results

PLK2 is reported to be responsible for phosphorylation of α-syn in murine brain and may potentially be responsible for the pS129 phosphorylation of the α-syn aggregates abundant in PD and other synucleinopathies. To investigate the role of PLK2 in this process, we used an organotypic brain slice model where aggregation, pS129 phosphorylatation and spreading of α-syn pathology is induced via injection of α-syn S129A-PFF [13], [33] by using pharmacological inhibition of PLK2.

### PLK2 inhibition reduces physiological nuclear pS129 levels but does not affect S129-phosphorylation of PFF-induced α-syn aggregates or their formation

To assess the effect of PLK2 inhibition on PFF-seeded aggregation using OHSCs, slices were treated continuously with 20 μM PLK2i (compound 37), from 24 hours prior to S129A PFFs injection until the end of experiment. PLK2i was added at each change of medium to ensure continuous inhibition of PLK2 function during the experiment (Fig. 1a). Seven days following S129A PFF-injection, slices were collected for biochemical analysis and immunostaining, at which time point the organotypic slice cultures normally express robust levels of total and pS129 α-syn (Fig. 1b). This time point allows the templating of α-syn aggregates in neurons at the site of injection at DG and the spreading of α-syn aggregate pathology to neurons in the CA3 and CA1 regions as described earlier and the use of S129A PFF can ensure that the signals detected using antibodies against pS129 is detecting only the de novo generated aggregates in the slices and not detecting the injected PFF materials as described earlier in [13].

**Fig. 1:**
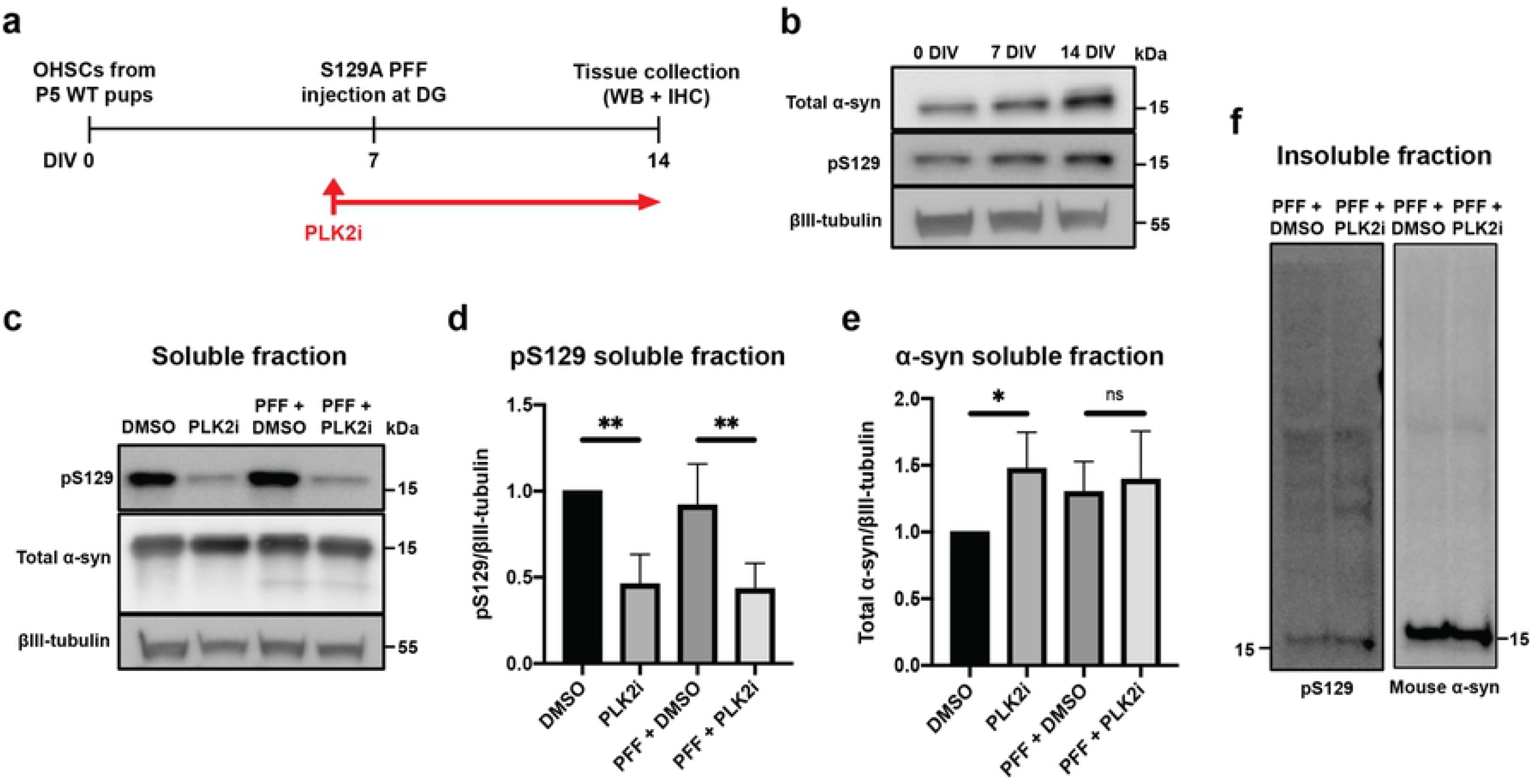
PLK2i treatment reduces S129-phosphorylated α-syn in the soluble fraction but not the insoluble fraction of organotypic slices. a) Workflow of the experiment. PLK2 inhibition is started 24 hours before S129A PFF injection and continued until tissue collection at 7 dpi. The timeline is depicted as days in culture (DIV). b) Immunoblotting of OHSCs from wild type (C57BL/6) pups, showing expression of endogenous α-syn and pS129 α-syn at 0, 7 and 14 days in culture, demonstrating the presence of a basal level of physiological RIPA-soluble pS129 α-syn in the slices. c) Immunoblotting of the RIPA-soluble fraction of PLK2i- or DMSO-treated slices ± PFFs for pS129 and total α-syn, demonstrating reduction of pS129 in PLK2i-treated slices in a PFF-independent manner. d & e) Quantification of immunoblots for pS129 (d) and total α-syn (e) in the RIPA-soluble fraction. Band values relative to βIII-tubulin were normalized to DMSO/non-injected group and represent mean ± SD, n = 3 independent experiments of 8 to 10 slices per group in each experiment. Statistical comparisons were performed using one-way ANOVA with Fisher’s LSD as post hoc test, * p < 0.05, ** p < 0.01. P-values for pS129 (d) were 0.004 (DMSO vs. PLK2i) and 0.007 (PFF+DMSO vs. PFF+PLK2i). P-values for total α-syn (e) were 0.0498 (DMSO vs. PLK2i) and 0.6607 (PFF+DMSO vs. PFF+PLK2i). f) Immunoblotting of the RIPA-insoluble fraction of PFF-injected slices ± PLK2i shows that PLK2 inhibition does not affect generation of aggregates, or S129-phosphorylation of α-syn in the insoluble fraction. Representative blot from 3 independent experiments. Molecular size markers in kDa are indicated to the right.

Immunoblotting of slice cultures at 7 days post injection (dpi) showed that PLK2i treatment increased the level of mouse α-syn in the soluble fraction (Fig. 1c & e) while the level of likely membrane associated pS129 in the RIPA-soluble fraction was reduced in a PFF-independent manner (Fig. 1c & d). By contrast, PLK2 inhibition did not affect the formation of insoluble α-syn aggregates as detected by immunoblotting using rodent-specific α-syn antibody D37A6 and pS129 extracted by 7M urea from the of RIPA-insoluble material of PFF-injected slice cultures (Fig. 1f).

IHC analysis of organotypic slice cultures confirmed the induction of aggregation of endogenous α-syn following S129A PFF-microinjection. Aggregates were detected by the aggregate-specific α-syn antibody, MJF-14, and pS129-α-syn (Fig. 2a). Staining for pS129 using the monoclonal antibody 11A5 revealed two forms of pS129 immunoreactive signals. One is bright and intense, aggregation-specific pS129-signals that co-localizes with MJF-14-signal and is only detected in PFF-injected slices (Fig. 2a, arrows). The other type is a fainter, diffuse non-aggregate specific pS129-signal, which is abundant in nuclei and co-localizes with DAPI-signal but never with MJF-14-signal, and is detected in both PFF and PBS-injected slices (Fig. 2a, arrowheads) but not in the PLK2i treated slices. This indicates that PLK2i-treatment significantly reduced the diffuse, nuclear pS129-signal in both PFF- and PBS-injected slice cultures (Fig. 2a, e). In contrast, PLK2 inhibition did not influence PFF-induced α-syn aggregation, as detected with aggregate specific MJF-14 antibody (Fig. 2b, c), and it did not prevent or reduce S129-phosphorylation of the aggregates generated in axons (Fig. 2d). The lack of dissociation between MJF-14 and pS129-signals in the PFF-induced aggregates following the PLK2i treatment demonstrates that aggregate-associated pS129 is unaffected by PLK2 inhibition (Fig. 2). Quantification analysis revealed an approximately 30% reduction of nuclear pS129 following PLK2i-treatment without reducing aggregate-specific pS129 inside MJF-14 signals (Fig. 2d & e). This led to an approximate doubling of the signal-to-noise ratio of aggregate-specific pS129 signal, thereby facilitating the study of pS129 positive α-syn aggregates without interference of non-aggregate-specific pS129 (Fig. 2f).

**Fig. 2:**
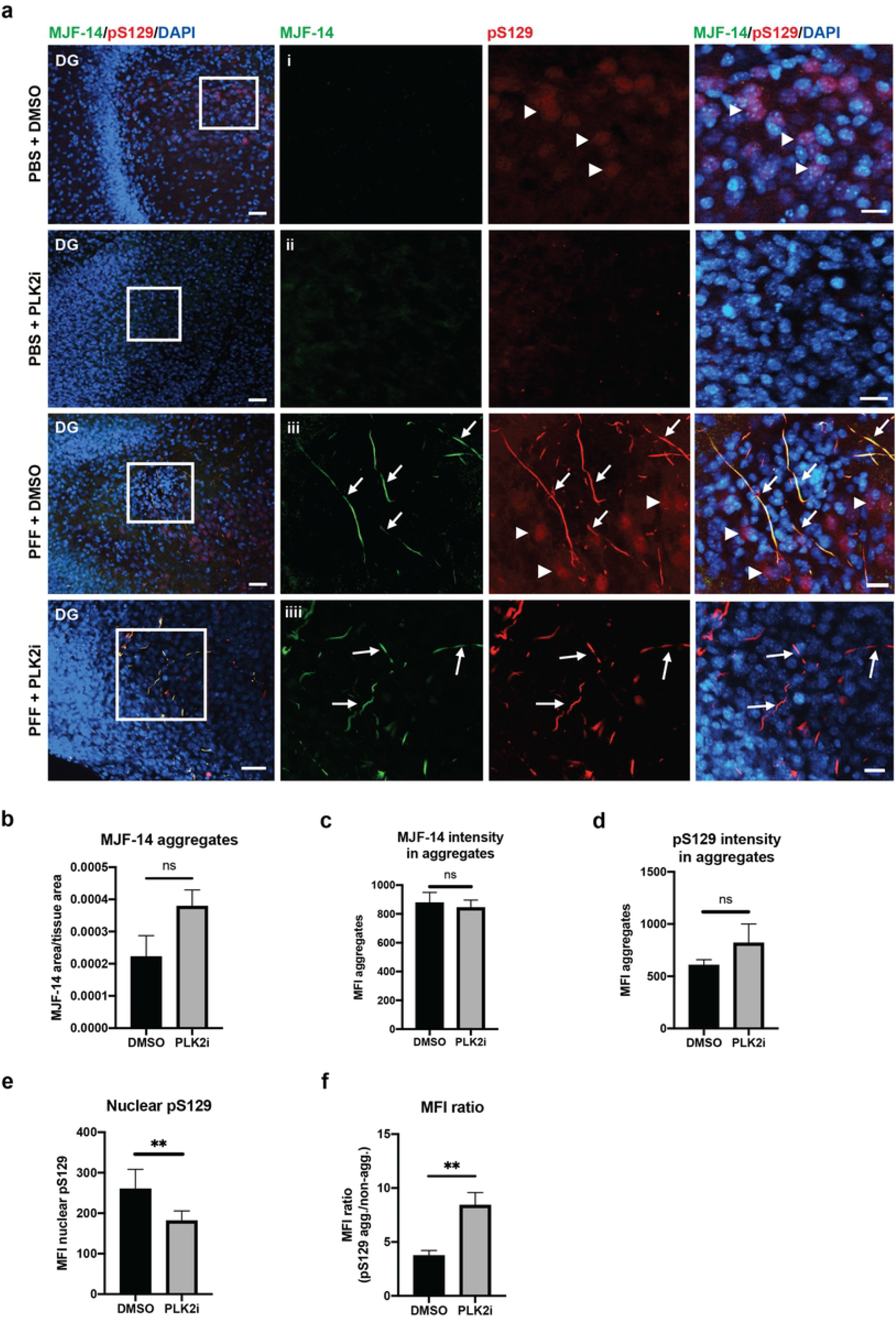
PLK2i treatment of OHSCs does not affect generation of pS129 positive aggregates, but reduces the nuclear pS129 intensity. a) Representative images from the dentate gyrus region (DG) of organotypic slices injected with PBS or S129A PFFs and treated with DMSO or PLK2i. Slices were stained with MJF-14, pS129 (11A5), and DAPI, scale bar = 50 μm. i. The magnified region (white boxed area) shows nuclear pS129 signal that co-localizes with DAPI (arrowheads). ii. The nuclear pS129-staining was removed following PLK2i treatment. iii. S129A PFF-induced α-syn aggregates, phosphorylated at S129, are detected with MJF-14 and pS129 antibodies (arrows). pS129-staining of non-aggregated α-syn is also seen, predominantly located in the nuclei (arrowheads). iiii. PLK2i treatment shows no effect on the S129-phosphorylation of aggregated α-syn (arrows) but effectively reduces the nuclear pS129-staining. Scale bars = 20 μm. b) Quantification of the amount of aggregation (aggregate area normalized to tissue area), defined by MJF-14 staining (p-value = 0.09). c-d) Quantification of the mean fluorescence intensity of α-syn aggregates detected with MJF-14 (c, p-value = 0.712) and aggregate-specific pS129 that overlaps with MJF-14 (d, p-value = 0.214). e) Quantification of nuclear (non-aggregate related) pS129, illustrating a significant decrease in mean fluorescence intensity following PLK2i-treatment (p-value = 0.0087). f) Relative ratio of aggregate-specific pS129/nuclear pS129 in PFF-injected slices shows an increase of ratio of the signals of aggregates/background following PLK2i treatment, due to reduction of the non-aggregate-related nuclear pS129-signal (p-value = 0.009). Bars represent mean ± SD of 3 independent experiments with 8 to 10 slices/experiment. Treatments were compared using an unpaired Student’s T-test.

### PLK2 inhibition does not hinder inter-neuronal spreading of PFF-induced α-syn aggregates

To test the influence of PLK2 inhibition on the spreading of PFF-induced α-syn aggregate pathology, aggregate signals at CA1 region was identified for quantification (Fig. 3a). PLK2i-treatment of slices showed no significant influence on aggregates formed at CA1 following the PFF injection in the DG when compared to DMSO-treated slices and their phosphorylation as determined by aggregate- and pS129-specific antibodies (Fig. 3b, arrow), but instead only reduced the nuclear pS129 signals (arrow head) (Fig. 3b-d). A tendency towards increased pathology load was observed, as also seen in Fig. 2b, although this was not statistically significant. To obtain a measure for the relative inter-neuronal spreading in each slice, the aggregate signal at CA1 was normalized to the aggregate signal at DG (Fig. 3e).

**Fig. 3:**
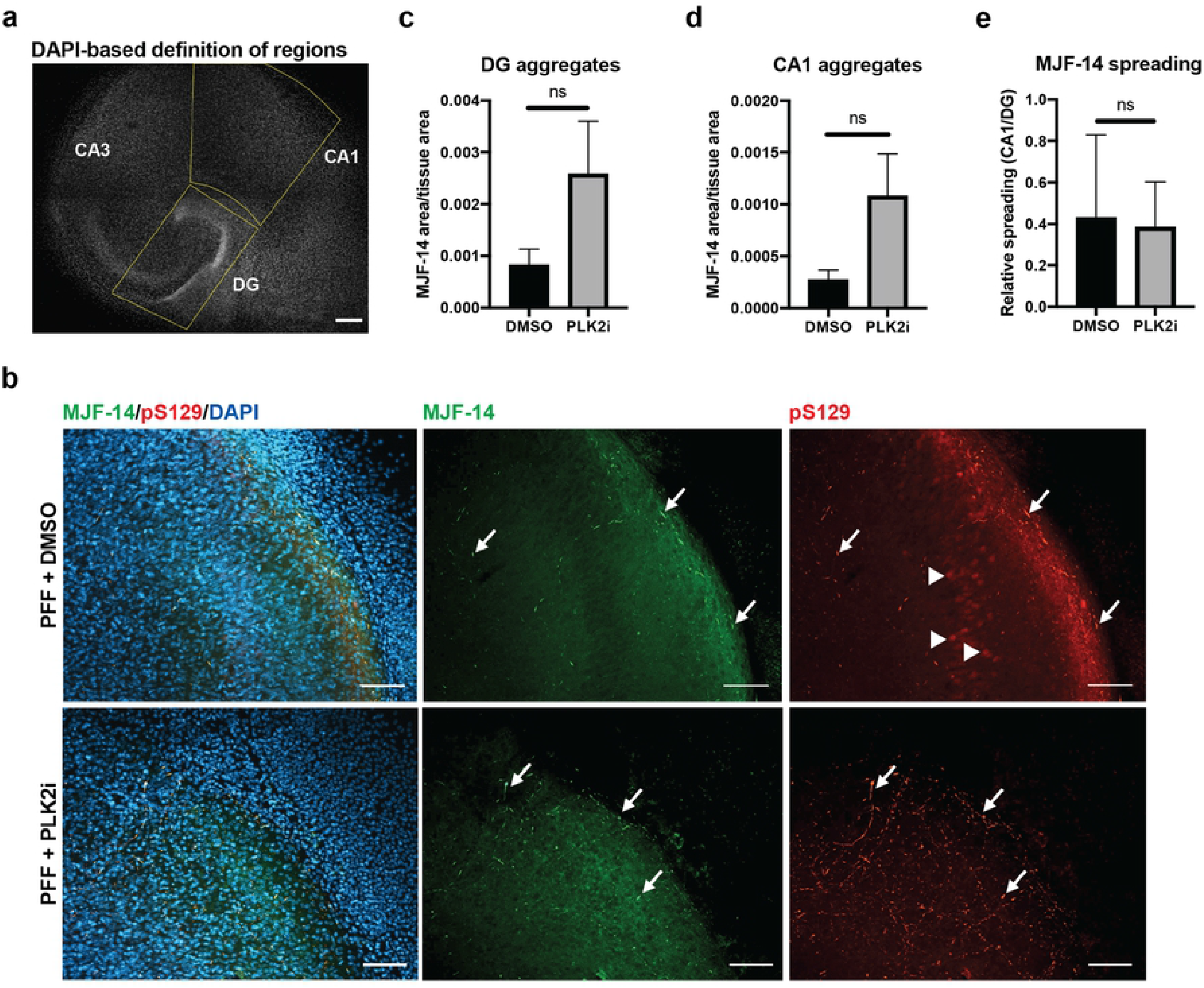
Inter-neuronal spreading of a-syn aggregate pathology from the DG to the CA1 region occurs independently of PLK2 inhibition. a) Segmentation of DG and CA1 regions in organotypic hippocampal slices labelled with DAPI, shown in gray scale. The DG is bounded manually by a box, and the CA1 regions is afterwards defined by extrapolation of one side of the DG box and a diagonal line through the box. Scale bar = 200 μm. b) Immunostaining of S129A PFF-injected OHSCs showing aggregated α-syn at the CA1 region 7 dpi detected by both MJF-14 and pS129. PLK2i-treatment reduces the nuclear pS129-signal at CA1 that overlaps with DAPI (arrowheads) but has no influence on the PFF-induced aggregate-specific pS129-signal that overlaps with MJF-14-signal (arrows). Scale bars = 50 μm. c-d) Quantification of α-syn aggregate area (MJF-14 area normalized to tissue area) at the DG (c) and CA1 region (d) shows no significant effects on aggregation at both regions following PLK2i treatment. e) Aggregate levels at the CA1 region were normalized to aggregate levels in DG of the same slice to address the relative spreading of S129A PFF-induced α-syn aggregates, based on MJF-14 staining. PLK2i treatment did not affect relative spreading of MJF14-positive pathology (p-value = 0.829 using an unpaired Welch’s T test). Bars represent mean ± SD of 5-6 slices per group. Images are representative of three independent experiments.

As PLK2 inhibition did not affect either the PFF induced aggregation, S129 phosphorylation or spreading of α-syn aggregates, but only diminished the diffuse nuclear pS129-signal in our slice cultures setup, we tested whether short-term treatment of slices with PLK2i would reveal the same result. Treatment of OHSCs with PLK2i or the PLK1-3 inhibitor (BI2536) for 24 hours prior to fixation reduced nuclear pS129 intensity, with a concomitant increase of the signal-to-noise ratio of aggregate-specific pS129-signal similar to the continuous PLK2 inhibition during the culture period (Fig. 4).

**Fig. 4:**
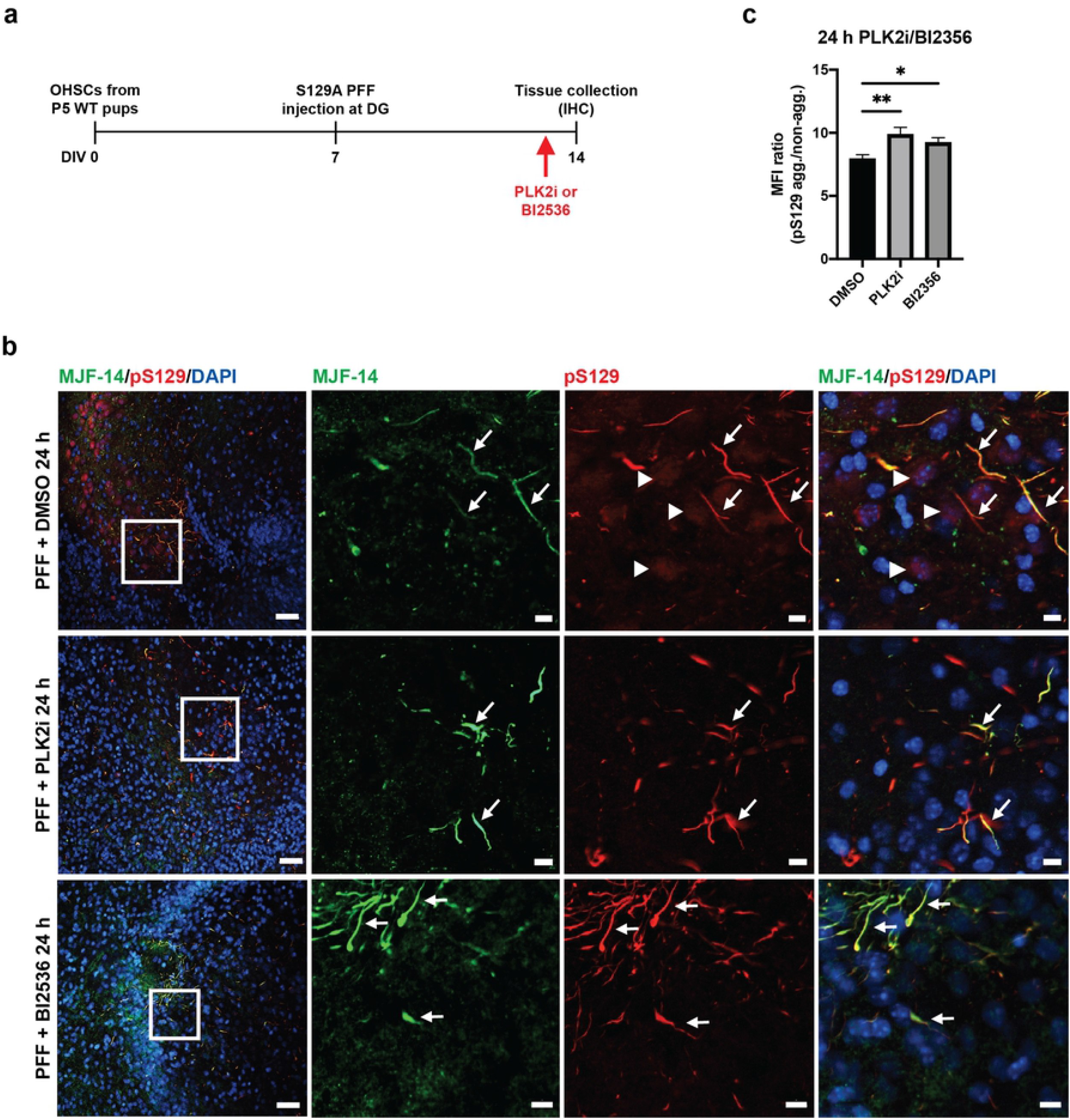
24 hours treatment of OHSCs with either polo-like kinase inhibitor reduces the nuclear pS129-signal. a) Experimental setup, with addition of PLK2i or PLK1-3 inhibitor (BI2536) to the medium 24 hours prior to slice collection. b) Immunostaining with MJF-14, pS129 (11A5) and DAPI, shows the reduction of nuclear pS129-staining (arrowheads) in PLK2i and BI2536 treated slices, while aggregate-specific pS129-staining that colocalizes with MJF-14-signals remains. Scale bars = 50 μm for the left column and 20 μm for the magnified inserts in three rightmost columns. c) Quantification showing an increase in the ratio of mean fluorescence intensity between aggregate-specific pS129 and nuclear pS129 signal following treatment with PLK2i (p-value = 0.0039) or BI2536 (p-value = 0.0467). No difference was observed between PLK2i and BI2536 (p-value = 0.3060). Graph displays mean ± SD of 6 – 8 slices per group. Statistical comparisons were performed using one-way ANOVA with Fisher’s LSD test, * p < 0.05, ** p < 0.01. Images are representative of three independent experiments.

### PLK2 inhibition reduces nuclear pS129 signal in the M83 model of synucleinopathy as well as in human neurons

To further substantiate the efficacy of a short-term PLK2 inhibition paradigm as a method to facilitate easier detection of α-syn aggregate pathology by the reduction of non-aggregate specific pS129 signals, we treated 15-month-old heterozygous A53T-α-syn transgenic mice (M83^+/-^) with PLK2i for 48 hours before sacrifice. Aggregate pathology in the models was initiated by injection of mouse recombinant PFFs in the hind limb gastrocnemius muscle at 12 months [29]. Immunostaining of the brain using anti-pS129 antibody showed a significant reduction in nuclear pS129 intensity as demonstrated in representative images from hippocampus and frontal cortex (Fig. 5a-c). The level of reduction varied between regions, with nuclear pS129 α-syn of pyramidal neurons of the CA1 region in the hippocampus appearing particularly sensitive to PLK2 inhibition (Fig. 5a, d). In contrast, the frontal cortex appeared to contain both PLK2i-sensitive and PLK2i-resistant nuclear pS129-signals (Fig. 5c, f). Quantification the pS129-positive aggregated fibrillary signals detected in the hind brain region showed that treatment of the mice with PLK2i for 48 hours – unsurprisingly – had no effect compared to the control group, considering the short time of the treatment (Supplementary fig. 3a & b). However, PLK2 inhibition strikingly increased the signal-to-noise ratio, facilitating the detection of aggregate-specific signals in the PLK2i treated mice (Supplementary fig. 3c).

**Fig. 5:**
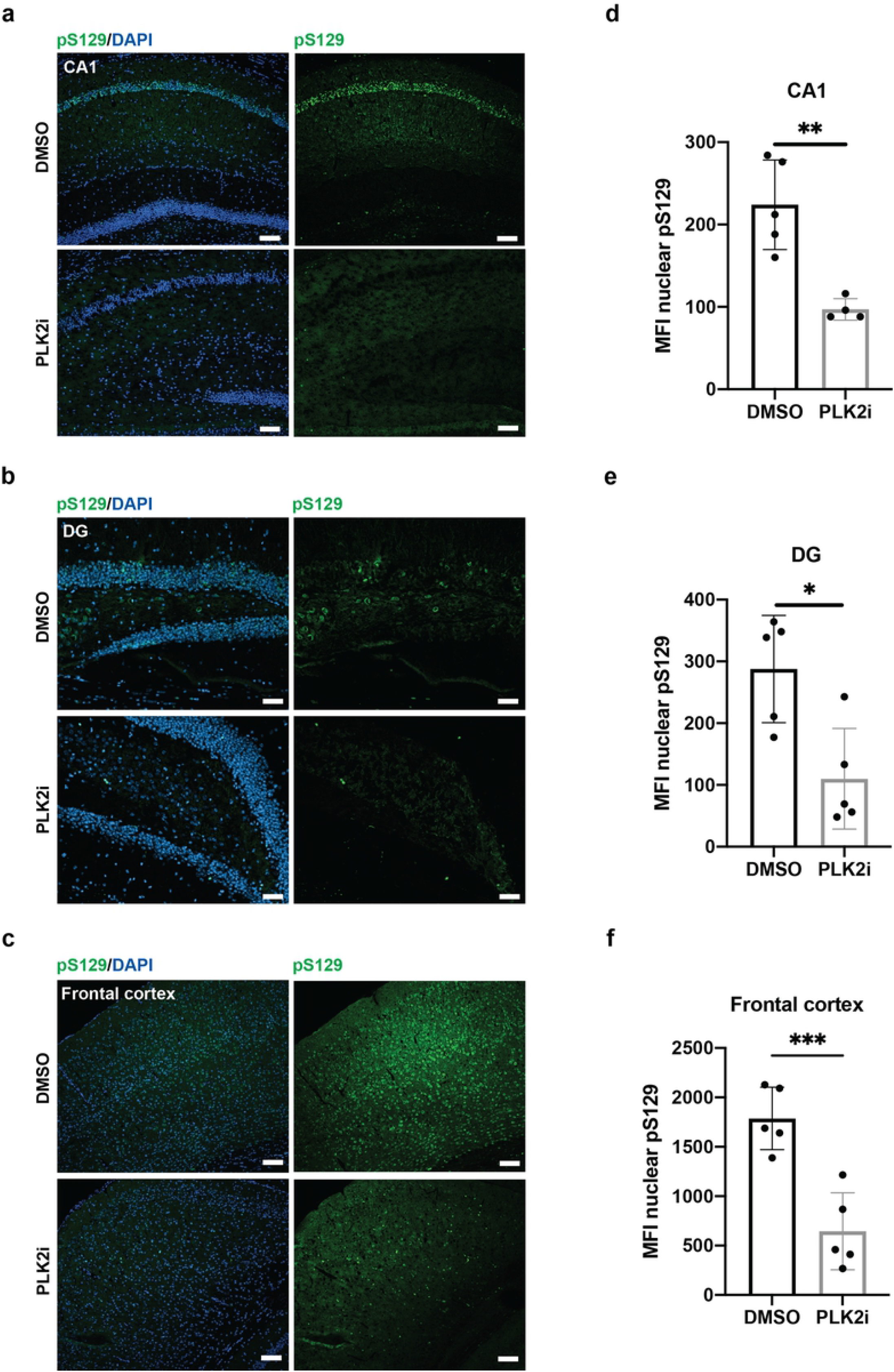
PLK2i treatment for 48 hours before sacrifice reduces nuclear pS129-staining in PFF-injected M83 mice. a-c) Representative immunostaining images from brain sections the of 15-month-old M83 mice – 3 months following PFF injection in the hindlimbs – stained with pS129 (D1R1R) and DAPI. CA1 region (a) and the DG (b) of hippocampus, and the frontal cortex (c), showing particularly strong pS129-staining at pyramidal neurons of the CA1 region. 48-hour PLK2i treatment by oral gavage (2 x 100 mg/kg/day) significantly reduces nuclear pS129 staining in all regions, although some PLK2i resistant S129-phosphorylation is apparent in the frontal cortex. Scale bars = 50 μm. d-f) Quantification of the mean fluorescence intensity of nuclear pS129-signal in CA1 (d, p-value = 0.0028), DG (e, p-value = 0.0102) and frontal cortex (f, p-value = 0.0009) demonstrate reduction of pS129 nuclear intensity following PLK2 inhibition. Bars represent the mean ± SD, n = 5 mice in each group. * p < 0.05, ** p < 0.01, as determined by an unpaired Student’s T test.

We also conducted a short-term PLK2 inhibition on cultured human dopaminergic neurons, where aggregation of endogenous α-syn was initiated by adding α-syn S129A PFFs (Fig. 6a). Addition of PLK1-3 inhibitor (BI2536) four hours before fixing the cells effectively reduced nuclear pS129 intensity, especially in the tyrosine hydroxylase-positive neurons, which displayed intense nuclear pS129-staining in DMSO-treated cultures (Fig. 6b & c). No modulation of aggregate levels following PLK2 inhibition was identified in the cultures treated with S129A PFFs (Fig. 6b & d). Collectively, these results highlight the ability of short-term PLK2 inhibition prior to fixation to improve the signal-to-noise ratio of aggregate-related pS129-signal across model systems from neuronal cultures to ex vivo tissue slices and in vivo models.

**Fig. 6:**
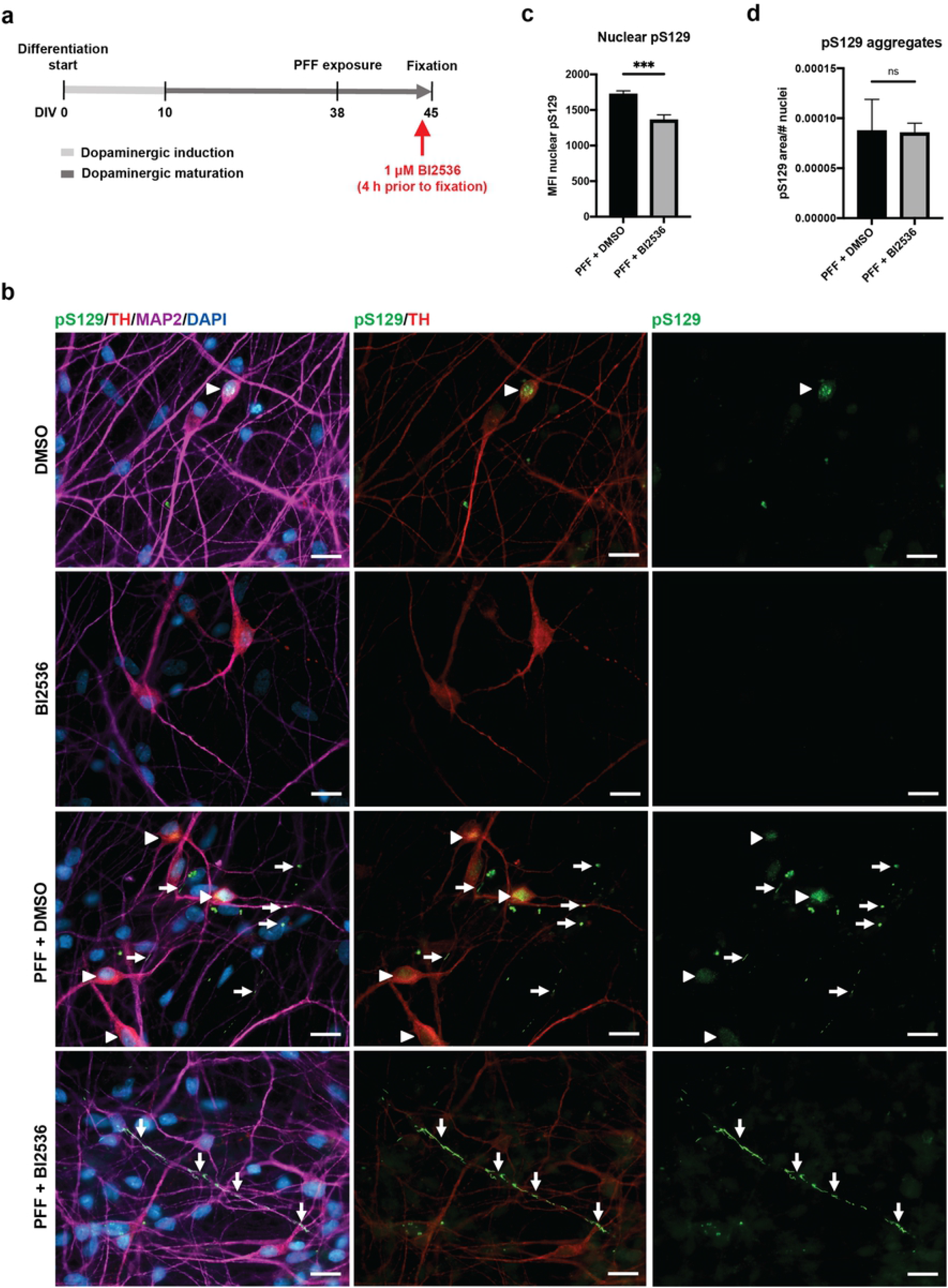
Inhibition of PLK2 reduces the nuclear pS129-signal in human dopaminergic neurons derived from iPSCs. a) Workflow of the experiment where the PLK1-3 inhibitor (BI2536) was added 4 h before fixing the cells for ICC. b) Representative images of iPSC-derived mature neurons visualized using MAP2 as a pan-neuronal marker, TH as dopaminergic neuronal marker, pS129 (11A5) and DAPI as a nuclear counterstain. For DMSO-treated cells, the staining shows bright nuclear pS129-signals (arrowheads), which disappear when cells are treated with BI2536, leaving the aggregate-specific pS129-staining easily detectable (arrows). Scale bars = 20 μm. c) Quantification of the average nuclear pS129 intensity in DMSO- and BI2536-treated cultures after exposure to PFFs shows decreased intensity in the BI2536 group (p-value = 0.0002). A minimum of 4000 cells were analyzed per group. Data are shown as mean ± SEM, and significance is indicated as *** p<0.001 using an unpaired Welch’s T test. d) Quantification of pS129 aggregate area normalized to the number of nuclei shows no difference between DMSO- and BI2536-treated cultures after exposure to PFFs (p-value = 0.5079). Data are shown as mean ± SEM, and significance is tested with a Mann-Whitney test.

To further explore how short a period of PLK2 inhibition is needed to reduce nuclear pS129 levels, we tested PLK inhibition from 10 minutes to 4 hours prior to fixation and analysis in primary hippocampal neurons (Supplementary fig. 4a). As little as 30 minutes treatment with 10 nM BI2536 prior to fixation was sufficient to decrease nuclear pS129 staining to a minimal level, which remained unchanged at 2 and 4 hours of treatment (Supplementary fig. 4b & c). As indicated in Supplementary fig. 4b, the nuclear pS129 staining is not removed completely upon treatment of the neuronal culture using BI2536, but it facilitates the study of the pS129-positive aggregates by increasing the signal-to-noise ratio as demonstrated in our cell, tissue and in vivo models.

## Discussion

The phosphorylation of α-syn at serine 129 has for almost two decades been recognized as a prominent characteristic of pathological α-syn aggregates not only in human tissue but also in cell and animal models of α-syn aggregate pathology [3],[4],[34],[35],[36],[37],[38]. However, the exact role of S129-phosphorylation remains contested due to conflicting results from various cell and in vivo studies [6], [7], [8], [9], [10], [11], [12],[13]. Those conflicting results are believed to be due to the use of models were either α-syn or the studied kinases are over expressed or expressed in a mutated forms, including our own earlier report where mutated form of α-syn (S129G) that cannot be phosphorylated at S129 residue were expressed using viral vectors in organotypic slices made from SNCA genes knock out pups [13].

In the current paper, we investigate whether PLK2, which has been identified as an efficient S129-directed kinase in rodent brains [4],[17],[19],[20],[22], is truly a kinase responsible for S129-phosphorylation of α-syn aggregates, and if we can modulate the PFF induced α-syn pathology and the associated pS129 level by pharmacological inhibition of PLK2 using a set up that expresses both α-syn and PLK2 in a physiological levels.

As inhibitors, the specific PLK2 inhibitor, compound 37 [21],[23],[24], which has been validated earlier in our lab [23] and the PLK1-3 inhibitor BI2536 were used [17],[25], [39], [40]. Both are efficient inhibitors of PLK2 but whereas compound 37 is highly specific and well tolerated, it has to be custom synthesized. BI2536, on the other hand, is commercially available but reported to be toxic to mitotic cells [8], [40].

The low toxicity of compound 37 allowed us to treat the PFF-seeded organotypic slice cultures throughout the duration of the whole experiment. This demonstrated that inhibition of PLK2 does not affect the accumulation of aggregate-specific pS129. In contrast, PLK2 inhibition reduced the RIPA-soluble fraction of pS129 and elevated the level of soluble α-syn, corroborating earlier data reported where PLK2 inhibition led to increased α-syn level by increasing its transcription [23]. Our data are in line with a recently published study using GFP-α-syn mice crossed with PLK2 knock-out mice and other study from the same group that opted for pharmacological inhibition of PLK2 in zebra fish, demonstrating that PLK2 deletion or inhibition had no influence on S129-phosphorylation and aggregates formation [41], [42].

Prompted by the apparent potential of PLK2 inhibition to facilitate pS129 aggregate detection in immunostainings, we tested the effect of short-term PLK2 inhibitor treatment paradigms in various PFF-seeded models. In organotypic slices, identical results were obtained with short-term treatment of slices for 24 hours using either compound 37 or BI2536. The treatment with PLK2i and BI2536 was equally able to reduce nuclear pS129 staining in M83^+/-^ and human neuronal models following treatment for 48 hours and 4 hours prior to tissue or cell collection, respectively. In none of the models did the short-term PLK2 inhibition affect the phosphorylation of aggregated α-syn.

Interestingly, PLK2i treatment of M83 ^+/-^ model demonstrated a striking regional variance in the reduction of nuclear pS129. This could be explained by differences in the expression level of PLK2 in various neuronal subtypes in different brain regions of mice [43],[44], [45] and is corroborated by the varying effect of PLK2i on total pS129 levels in different parts of the mouse brain [22].

As the detection of pS129 in inclusions is often used as a readout to quantify aggregate formation and the effect of strategies directed to counteract this process [34], [36],[37], [38], [46], the presence of non-aggregate-associated pS129-signals represents a confounding signal that requires segmentation of immunofluorescence images. Based on our results, we propose short-term treatment with a specific PLK2i or BI2536 as an easy and efficient strategy to improve signal-to-noise ratios when quantifying aggregate-associated pS129 signals. A time-course analysis of BI2536-treated hippocampal neurons demonstrated efficient reduction in nuclear pS129 after as little as 30 minutes of treatment (Supplementary fig. 4). This timeframe is well in line with previous studies of the post-mortem stability of physiological pS129, where complete dephosphorylation is observed within 30 minutes of sacrifice, at which time point the kinase/phosphatase equilibrium is shifted towards dephosphorylation [3]. In contrast, aggregate-specific pS129 is a stable modification, prevalent in post-mortem patient brains many hours after death [3],[4].

Naturally, attention must be paid to other effects of the short-term PLK2 inhibition that was not tested here, but if the treatment is brief and the primary read-out is α-syn aggregates, which are considered fairly stable structures as demonstrated in the tested models in this study, then this concern should be of minor importance.

The presence of S129-phosphorylated α-syn in the nucleus has been questioned due to the reported off-target and non-specific binding of commercially available α-syn and pS129 antibodies [47], [48], [49], [50]. Nonetheless, others have validated the presence of soluble endogenous pS129 α-syn in the nucleus using antibodies with a high affinity to pS129 epitopes or using combination of different pS129 antibodies [20],[49],[51], [52]. With this in mind, the validity of the pS129-staining detected with the 11A5 antibody used in this study [4] was tested using slices from α-syn knock-out (ASKO) pups that do not express α-syn. Immunostaining of ASKO slices showed no off-target signals (Supplementary fig. 2) in accordance with our earlier data obtained by western blotting using 11A5 in ASKO slices [33], substantiating the specificity of 11A5 antibody and corroborating a previous validation of nuclear pS129. Our finding of physiological, non-pathology related pS129 α-syn is consistent with numerous studies of both in vitro and in vivo models, reporting the nuclear localization. Although our data do not identify the roles of nuclear pS129 α-syn, they provide a novel experimental strategy to investigate this enigmatic α-syn species that has been associated to processes covering histone acetylation, neurotoxicity, transcriptional regulation and repair of double-strand DNA breaks [6], [20], [51], [53], [54], [55], [56], [57].

### Conclusion

The findings of this study demonstrate that PLK2 is involved in significant S129-phosphorylation of physiological α-syn but not the phosphorylation of serine-129 on aggregated α-syn. Moreover, short-term PLK2 inhibition can be used as an easy experimental procedure to facilitate specific detection of aggregated α-syn in different models of templated α-syn pathology.

## Supplementary figures

**Supplementary fig. 1: Characterization of the mouse recombinant PFFs injected in M83 mice**. a) Biochemical characterization of mouse PFFs. The insoluble fibrils consist of pure α-syn as demonstrated by SDS-PAGE and Coomassie blue staining (P = pellet, S = supernatant). b) The sonicated mouse PFFs comprise a homogeneous, mono-dispersed particle population with a 38.8 nm radius as determined by dynamic light scattering (DLS). c) The amyloid nature of the PFFs was confirmed by a robust K114 fluorometric signal detected at 550 nm. In comparison, monomeric α-syn did not produce any signal.

**Supplementary fig. 2: Validation of pS129 antibody (11A5) for immunostaining.** In OHSCs made from α-syn knock out (ASKO) pups and injected with S129A PFFs, immunostaining using pS129 (11A5) yields no signal at 7 dpi. In comparison, slices from WT (C57BL/6) pups injected with S129A PFFs demonstrate a nuclear pS129-signal that co-localizes with DAPI signals, predominantly in the pyramidal neurons of the CA3 and CA1 region of hippocampus (arrowheads). The nuclear signals are more diffuse and less bright than the PFF-induced axonal α-syn aggregate signals (arrows). Scale bar = 100 μm. i and ii. Magnified inserts show the bright distinct PFF-induced aggregates (arrows) and more diffuse non-aggregate-specific nuclear pS129 signal (arrowheads). Scale bars = 20 μm.

**Supplementary fig. 3: 48 hours PLK2i treatment of M83 mice has no influence on the generated α-syn aggregates**. a) Representative images of α-syn aggregates detected in the hind brain of M83 mice, scale bar = 50 μm. b) Quantification of the pS129 aggregate area normalized to tissue area. Aggregate amount is unaffected by PLK2i treatment (p-value = 0.3075 using an unpaired Welch’s T test). c) PLK2i treatment facilitates easier detection of pS129-positive aggregates, as the ratio of mean fluorescence intensity of pS129 between aggregates and non-aggregate nuclear signal increases drastically upon treatment (p-value < 0.0001 by a Mann-Whitney test). Bars represent the mean ± SD, n = 5 mice per group.

**Supplementary fig. 4: PLK2 inhibition reduces the intensity of nuclear pS129 staining in as little as 30 minutes.** a) Experimental overview for the determination of PLK2 inhibition time course in primary hippocampal neurons. b) Representative images from hippocampal neurons cultured for 14 days and treated with 10 nM BI2536 for 0-4 hours prior to fixation. A minimum of 30 minutes treatment was sufficient to effectively decrease nuclear pS129 but not remove it completely, as is demonstrated by the increased exposure images on the right. Scale bars = 20 μm. c) Quantification of nuclear pS129-staining shows a plateauing of mean fluorescence intensity after 30 minutes of BI2536 (p-value = 0.0185). No decrease in nuclear pS129 was detected with 10 minutes treatment (p-value = 0.4028). Bars represent mean ±SD from 2 independent replicates and significance is indicated as * p<0.05 by one-way ANOVA followed by Holm-Šidák posttest.

